# Deconvolution of cellular subsets in human tissue based on targeted DNA methylation analysis at individual CpG sites

**DOI:** 10.1101/2020.07.28.225185

**Authors:** Marco Schmidt, Tiago Maié, Edgar Dahl, Ivan G. Costa, Wolfgang Wagner

## Abstract

**Background:** The complex composition of different cell types within a tissue can be estimated by deconvolution of omics datasets. For example, DNA methylation (DNAm) profiles have been used to establish an atlas for multiple human tissues and cell types. In this study, we investigated if deconvolution is also feasible with individual cell-type-specific CG dinucleotides (CpG sites), which can be addressed by targeted analysis, such as pyrosequencing.

**Results:** We compiled and curated a dataset of 579 samples from Illumina 450k BeadChip technology that comprised 14 different purified and characterized human cell types. A training and validation strategy was applied to identify and test cell-type-specific CpGs. Initially, the amount of fibroblasts was estimated using two CpGs that were either hypermethylated or hypomethylated in fibroblasts. This FibroScore correlated with the state of fibrosis and was associated with overall survival in various types of cancer. Furthermore, we identified hypomethylated CpGs for leukocytes, endothelial cells, epithelial cells, hepatocytes, glia, neurons, fibroblasts and induced pluripotent stem cells. Using previously published BeadChip datasets with cell mixtures the accuracy of this eight CpG signature was comparable to previously published signatures based on several thousand CpGs. Finally, we established and validated pyrosequencing assays for the relevant CpGs that can be utilized for classification and deconvolution of cell types.

**Conclusion:** This proof of concept study demonstrates that DNAm analysis at individual CpGs reflects the cellular composition of cellular mixtures and different tissues. Targeted analysis of these genomic regions facilitates robust methods for application in basic research and clinical settings.

## Background

The human body comprises hundreds of different cell types, but a clear and commonly accepted classification is still elusive (1). The cellular characterization is usually based on ontogenetic origin within a tissue, cellular morphology, and particularly on expression of cell-type-specific surface markers. These markers can also be used to isolate and purify distinct cellular subsets, e.g. by flow cytometry upon labeling with specific antibodies. However, most cell types do not have a unique panel of surface markers and bulk analysis without physical sorting masks the contribution of rare cell types (2, 3). In the advent of single cell omics data, e.g. by transcriptomics, ATAC-seq, or even single cell proteomics, it is possible to discern between cells by molecular means on a cell-by-cell basis (4). However, these methods require fresh material, they are relatively expensive, and clear demarcation of cell types remains a challenge. Alternatively, it is possible to use transcriptomic or epigenetic bulk data to estimate the cellular composition in tissues based on deconvolution algorithms (2, 5-8). Better insight into the composition of cell types may support pathological assessment, target identification, and staging of various diseases (6, 9). To this end, a robust, simple and cost effective method to estimate the cellular composition in a given tissue sample would be advantageous.

DNA methylation (DNAm) at CG dinucleotides (CpGs) is a stable and heritable modification that is directly linked to cellular differentiation (9-11). It can be analyzed quantitatively on single base resolution and - in contrast to gene expression - every cell has only two alleles, which makes DNAm ideally suited for deconvolution approaches (12). Amongst the first applications was the estimation of leukocyte subsets in blood (5). More recently, it has been shown, that comprehensive human cell type DNAm profiles facilitate the estimation of the origin of circulating cell-free DNA (7). Deconvolution may either be based on a reference dataset, or it can be trained reference-free (9, 13, 14). So far, epigenetic deconvolution was mostly based on genome wide DNAm profiles, generated by the Illumina BeadChip technology. This method is relatively cost-effective and provides a very broad insight into genome-wide DNAm patterns. However, targeted methods for DNAm analysis, such as pyrosequencing of specific CpGs, may facilitate faster and even more cost-efficient analysis with less starting material, while reducing batch to batch variation and other technical challenges (15). We have recently developed targeted DNAm signatures for pyrosequencing of individual CpGs to achieve deconvolution of leukocyte subsets that correlate with conventional blood counts (16). In this study, we followed the hypothesis, that the relative proportion of fibroblasts, or even the complex cellular composition of human tissues, can be estimated by targeted analysis of DNAm at individual CpGs.

## Results

### Compilation of global DNAm profiles of different cell types

To identify cell-type-specific CpGs for targeted methylation assays and tissue deconvolution, we curated and compiled 579 samples from 46 different studies, mostly generated with the Illumina 450K BeadChip technology. We only considered non-malignant samples and retrieved datasets of the following purified and characterized human cell types: fibroblasts, mesenchymal stromal cells (MSCs), adipocytes, astrocytes, leukocytes, endothelial cells, melanocytes, epithelial cells, glia, hepatocytes, muscle cells, muscle stem cells, neurons, and induced pluripotent stem cells (iPSCs). 409 samples were used as a training set and 170 samples from independent studies were used as a validation set (Supplemental Figure S1A; Supplemental Table S1). Multidimensional scaling (MDS) of genome wide DNAm profiles revealed that samples of the same cell type cluster together across different studies, supporting the notion that the cell type has major impact on DNAm patterns (Figure 1A; Supplemental Figure S1B).

**Figure 1:**
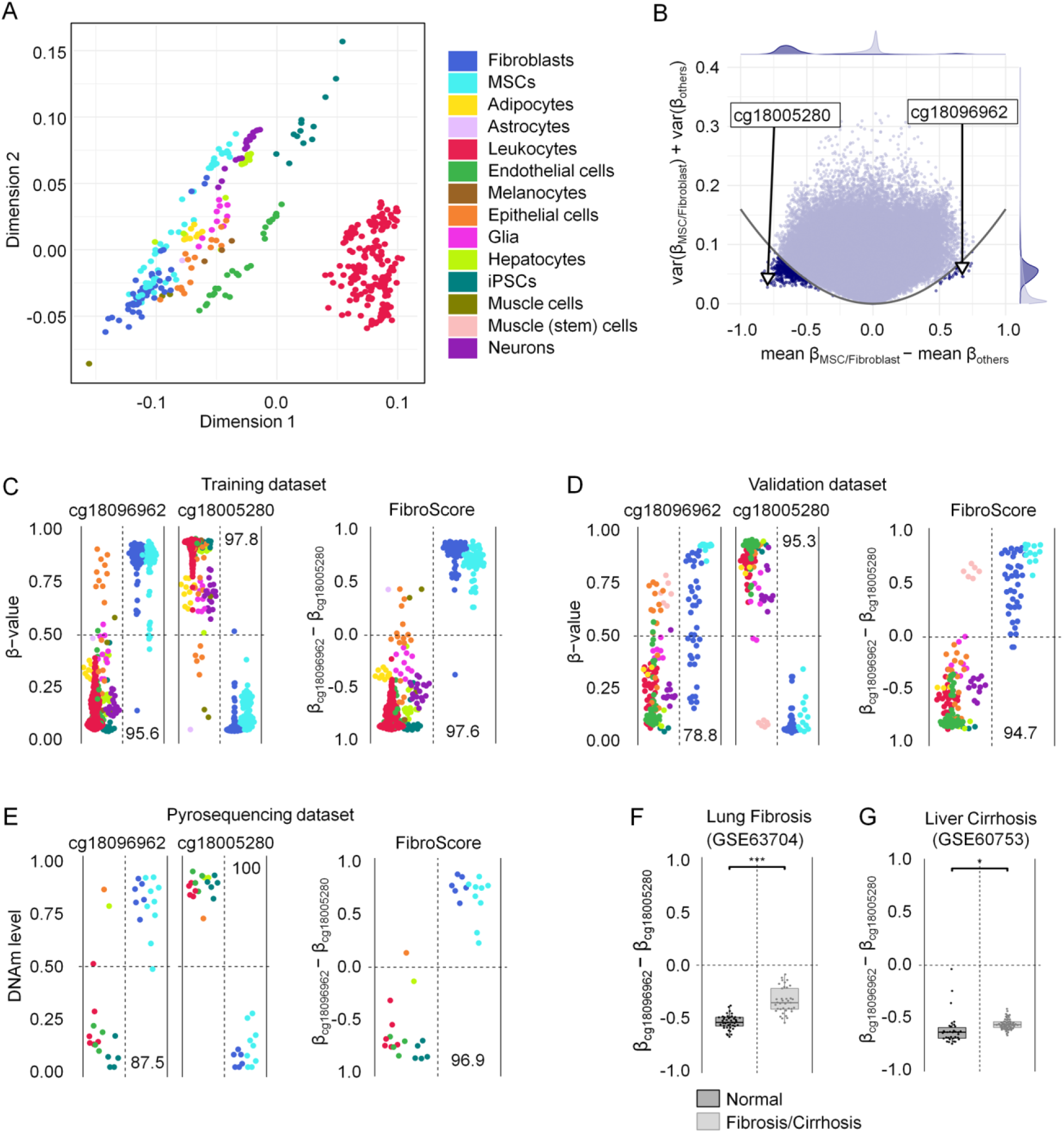
Selection of cell-type specific CpGs for fibroblasts. (A) Multidimensional scaling (MDS) plot of the training data set (n = 409) demonstrates that samples cluster by cell type across different studies. All CpGs shared between the 450K and EPIC BeadChip were considered (except XY chromosomes). (B) Differential mean DNAm levels of fibroblasts/MSCs *versus* all other cell types were plotted against the sum of variances within both groups. The CpGs, which have been selected for the FibroScore, are indicated. (C) DNAm levels (β values) of the two selected CpGs of the FibroScore in the training set. Numbers correspond to classification accuracy in percentage values. (D) DNAm levels of the two selected CpGs and the FibroScore for the validation set. Only muscle stem cells, which might closely resemble MSCs, were classified with fibroblasts/MSCs. Numbers correspond to classification accuracy in percentage values. (E) DNAm levels of the two selected CpGs and the FibroScore as determined by pyrosequencing in samples of different cell types. Almost all cell preparations (with exception of the HaCat cell line) were classified correctly. (F) The FibroScore is significantly higher in lung fibrosis *versus* healthy control tissue (GSE63704; 450K data)(19). *** p < 0.001. (G) The FibroScore is significantly higher in liver cirrhosis *versus* healthy control tissue (GSE60753; 450K data)(20). * p < 0.05.

### DNA methylation at fibroblast-associated CpGs can be indicative for fibrosis

To gain insight into the cellular composition of tissue, we initially selected CpGs that might discern fibroblasts from other cell types. Such fibroblast-specific DNAm patterns could reflect the relative proportion of fibroblasts, for example for staging of fibrotic diseases. In our previous work, we addressed differences in DNAm profiles of fibroblasts *versus* MSCs, albeit classification of these cell types is hardly reflected by clear functional or molecular characteristics (17). This is also reflected by their close relationship in the MDS plot. Therefore, we have decided to group both cell types together into the fibroblast category for subsequent analysis. To select fibroblast-specific CpGs that are either characteristically methylated or unmethylated in fibroblasts we filtered for CpGs based on (1) the highest difference in mean DNAm in fibroblasts *versus* other cells and (2) small variance in DNAm levels within each of the two groups (Figure 1B). CpG candidates were evaluated and ranked based on results of a 10-fold cross validation setup. Based on this we selected cg18096962 (associated with the lncRNA *RP11-60A8.1*) as hypermethylated and cg18005280 (associated with the gene leucine rich repeats and immunoglobulin like domains 1 [*LRIG1*]) as hypomethylated CpG site (Supplemental Figure S1C). The difference in DNAm levels between these CpGs ([β-value at cg18096962] – [β-value at cg18005280]), referred to as FibroScore, could clearly distinguish fibroblasts from most other cell types (Figure 1C,D). Only muscle stem cells, which have been differentiated for 24h toward myogenic lineage and might therefore closely resemble MSCs, were classified in the fibroblast category (18). To further validate applicability of these CpG sites for targeted DNAm analysis, we analyzed DNA samples from cultured cells, frozen blood, and commonly used cell lines with pyrosequencing (Figure 1E). Only one immortalized cell line was misclassified by the FibroScore: HaCat (spontaneously transformed keratinocytes for epithelial cells), which might be due to aberrant DNAm patterns by malignant transformation. Thus, targeted analysis of the two CpGs might be indicative for the fraction of fibroblasts/MSCs in tissue. In fact, when we applied the FibroScore to Illumina BeadChip datasets of lung fibrosis (GSE63704, Figure 1F; Supplemental Figure S1D) and liver cirrhosis (GSE60753, Figure 1G; Supplemental Figure S1E) we observed a significant higher FibroScore in the fibrotic tissues as compared to healthy controls (two-sided t-test: p = 2.51*10^−12^, and p = 0.0396, respectively).

### FibroScore correlates with overall survival in various types of cancer

Cancer-associated fibroblasts (CAFs) determine the tumor microenvironment and play a crucial role for progression of malignancies (21). Therefore, we anticipated that the FibroScore might also be of prognostic relevance for various types of cancers. To address this question, we utilized 32 datasets from The Cancer Genome Atlas (TCGA) and determined the FibroScore based on the DNAm at the two relevant CpGs. For each cancer type the patient data was then stratified by the median FibroScore. A higher FibroScore was indicative for significantly shorter overall survival in adrenocortical carcinoma (TCGA-ACC, p = 0.00034), chromophobe renal cell carcinoma (TCGA-KICH, p = 0.012), mesothelioma (TCGA-MESO, p = 0.0011), and head and neck squamous cell carcinoma (TCGA-HNSC, p = 0.005) (Figure 2). Despite the relatively low number of non-censored patients, it was also significant for pheochromocytoma and paraganglioma (TCGA-PCPG, p = 0.0064). For all other cancer types the stratification by the median FibroScore did not reveal significant association with overall survival (Supplemental Table S2). However, a significant association was observed if we only stratified by DNAm at cg18096962 for brain lower grade glioma (TCGA-LGG, p = 0.0113) and uterine carcinosarcoma (TCGA-UCS, p = 0.0179). Furthermore, if we stratified by cg18005280 (*LRIG1*) we also observed significant results for pancreatic adenocarcinoma (TCGA-PAAD, p = 0.0090), kidney renal clear cell carcinoma (TCGA-KIRC, p = 0.0103), uveal melanoma (TCGA-UVM, p = 0.0001) and kidney renal papillary cell carcinoma (TCGA-KIRP, p = 0.0455). These results support the notion, that the DNAm at fibroblast-associated CpGs might be indicative for the fraction of CAFs, which is relevant for progression of various types of cancer.

**Figure 2:**
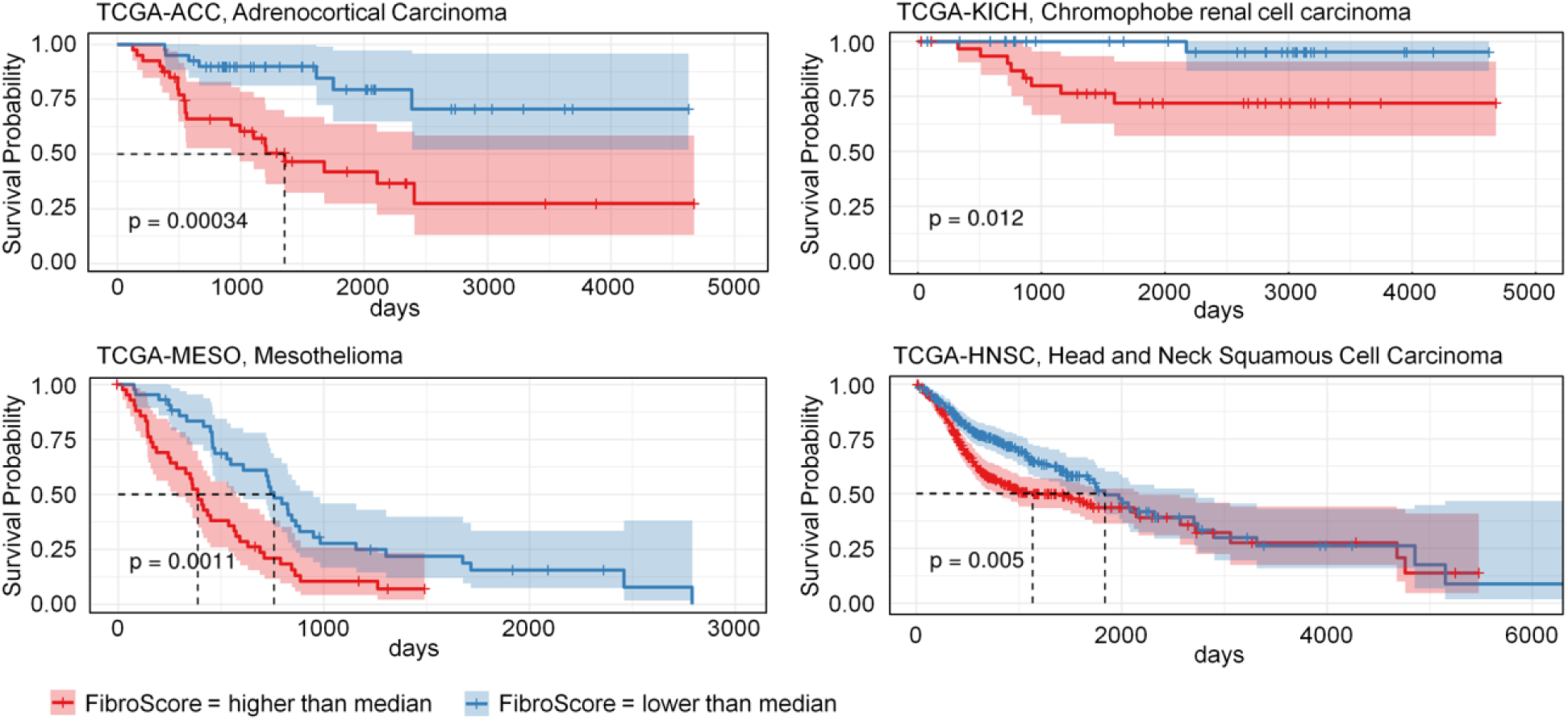
The FibroScore is associated with overall survival in several types of cancer. Kaplan-Meier curves of datasets from The Cancer Genome Atlas (TCGA) were stratified by the median FibroScore in 450K BeadChip data. Depicted are four types of cancer with significantly shorter overall survival in patients with higher FibroScore.

### Deconvolution of cell types based on individual cell-type specific CpGs

Subsequently, we followed the question if targeted analysis of individual CpGs might also reflect the composition of tissues. To this end, we have identified characteristic CpG sites for additional cell types using a similar procedure of CpG selection as mentioned above (difference in mean DNAm, variance in DNAm levels and classification performance). Notably, for all cell types – except for iPSCs, which resemble a ground state of non-differentiated cells – we identified more hypomethylated than hypermethylated CpGs in our feature selection (Figure 3A). One hypomethylated CpG site was selected for every cell type, most of which were within introns and exons of corresponding genes (Supplemental Figure S2): cg23068797 (*DNM2*, dynamin-2) for fibroblasts, cg10673833 (*MYO1G*, myosin IG) for blood cells, cg06631999 (*STMN1*, stathmin) for epithelial cells, cg27197524 (*POLE*, DNA polymerase epsilon catalytic subunit A) for hepatocytes, cg06421238 (*WSCD1*, WSC domain-containing protein 1) for endothelial cells, cg27309098 (*AGAP1*, Arf-GAP with GTPase, ANK repeat and PH domain-containing protein 1) for glia, and cg09998451 (*RAB3A*, ras-related protein Rab-3A) for neurons. Cell-type specific hypomethylation was validated with the Illumina BeadChip data from the validation set and by pyrosequencing of various cell types and tissues (Figure 3B).

**Figure 3:**
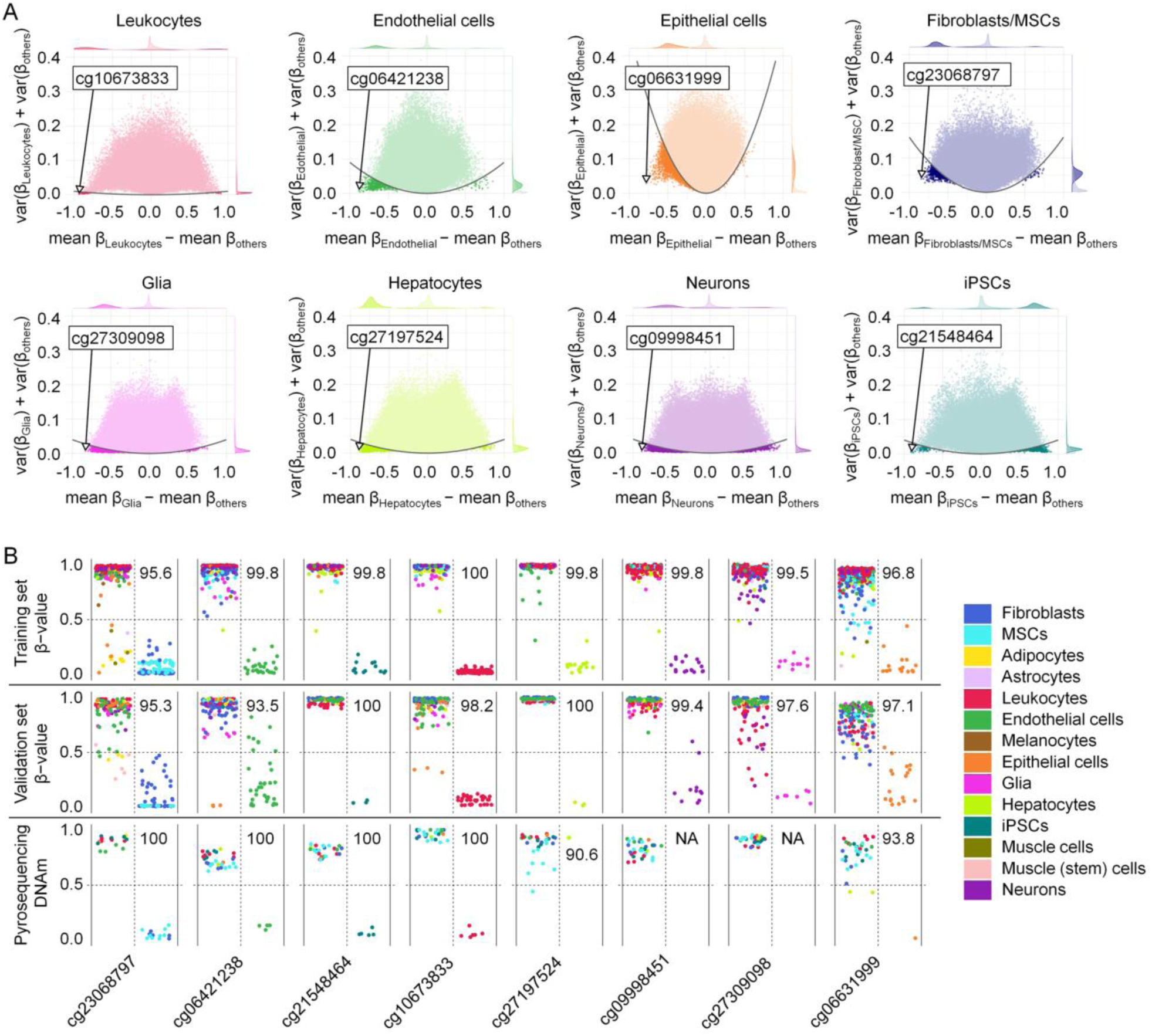
Cell-type-specific CpG sites are preferentially hypomethylated. (A) Selection of cell-type specific CpGs for leukocytes, endothelial cells, epithelial cells, fibroblasts/MSCs, glia, hepatocytes, neurons and iPSCs. The difference of mean β values of each cell type *versus* all other cell types was plotted against the sum of variances within both groups. CpGs for subsequent deconvolution are highlighted. (B) DNAm levels of the eight selected CpGs in the training, validation and pyrosequencing datasets. The vast majority of samples revealed the expected cell-type-specific hypomethylation, albeit pyrosequencing of liver cell lines (Hep3B and HuH-7) did not reveal hypomethylation at cg27197524 as expected for primary cells. Glia or neuron samples were not available for pyrosequencing. Numbers correspond to classification accuracy in percentage values.

We used the mean DNAm levels for the selected CpGs in eight distinct cell types in the training dataset as our reference matrix when applying the non-negative least squares (NNLS) deconvolution algorithm (Figure 4A; Supplemental Table S3). The NNLS algorithm could then be used to generate estimates for the cellular composition of tissues and DNA mixes, based on the DNAm of eight CpGs. To assess the performance of our deconvolution model we used a 450K Illumina BeadChip dataset of neuron-glia-DNA-mixes in incremental proportions (22). Our predictions correlated very well with the neuronal/glial proportions, with only a small fraction of other cell types being predicted as present (Figure 4B; Supplemental Figure S3A). Alternatively, we tested non-negative matrix factorization (NMF) and EpiDISH, which performed not as well as the NNLS approach (Supplemental Figure S3 A,B) (23, 24). Next, we tested the deconvolution performance on 450K data of DNA *in-vitro* mixes of various different cell types (7). Our training set did not include categories for lung and colon epithelial cells and therefore we assigned them to our epithelial cell category. Again the predictions of our NNLS approach overall closely represented the real composition of cell types, and the results were similar to the previously described deconvolution results that considered about 6000 CpGs (7) (Figure 4C; Supplemental Figure S3C). We then tested if deconvolution of different cell types would also be feasible by targeted methods. Therefore, we prepared five *in-vitro* DNA mixes of five different cell types in varying proportions and analyzed all cell-type-specific CpGs by pyrosequencing. The estimated composition closely resembled the previously mixed fractions (Figure 4D; Supplemental Figure S3D).

**Figure 4:**
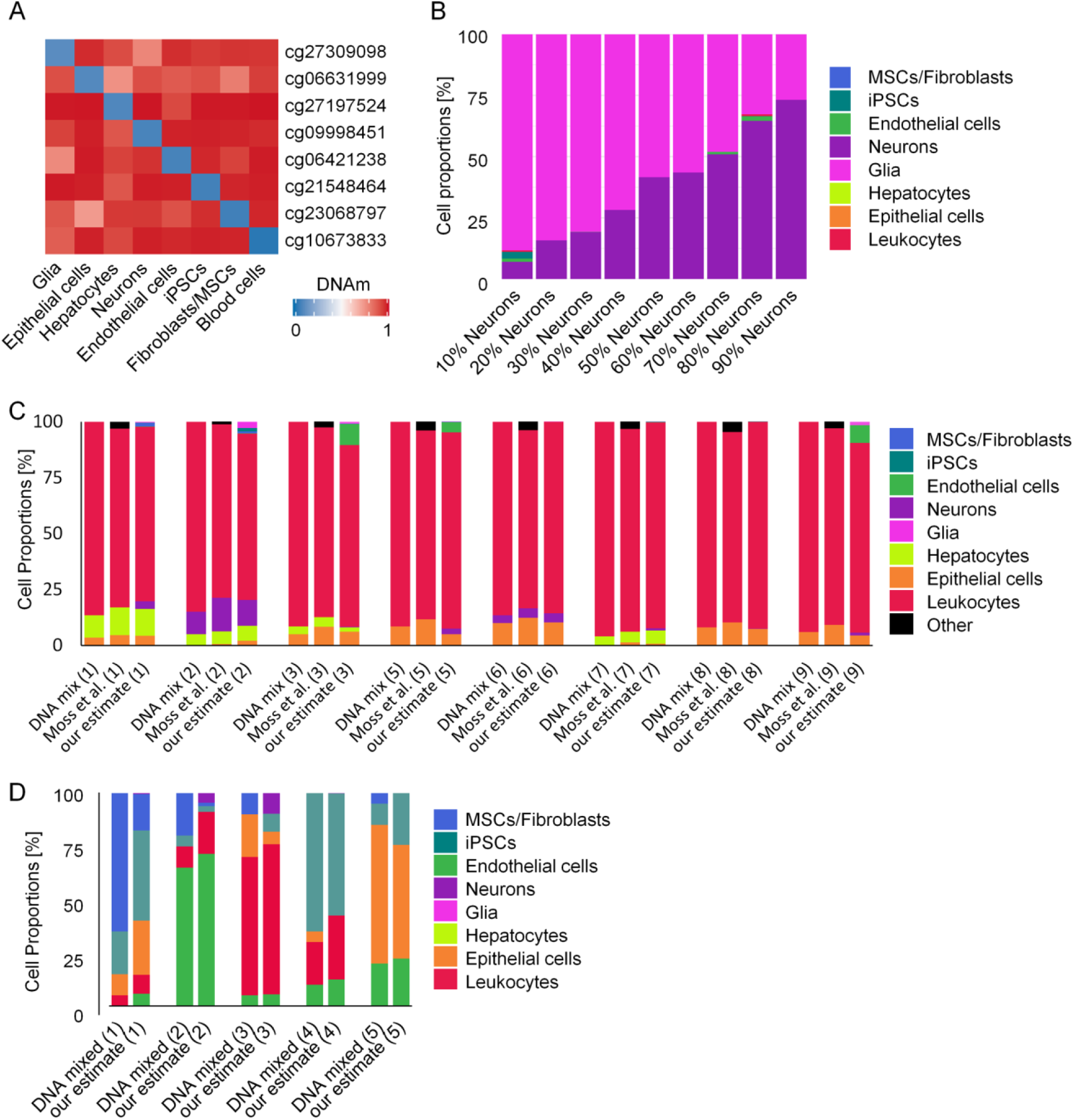
Deconvolution of cell mixtures based on individual cell-type specific CpGs. (A) Heatmap of mean β values of the reference matrix (450K data of the training set), which is used for deconvolution. (B) Deconvolution of *in vitro* neuron-glia-DNA-mixes from dataset GSE41826 (22). The predicted cell fractions by our NNLS-based deconvolution with eight CpGs are depicted. (C) Deconvolution of eight different *in vitro* DNA mixes from dataset GSE122126 (7). The real composition of DNA fractions is plotted next to the predictions by the signatures of Moss et al. (estimates for leukocyte subsets, epithelial cells and others were combined). The estimates with our NNLS model closely resembled the DNA mixtures of different cell types. Data of DNA mix 4 was lacking one of the eight CpGs and was therefore excluded. (D) Deconvolution of *in vitro* DNA mixes measured with pyrosequencing. Five different mixes of five different cell types in different proportions were measured at the eight different positions. Shown are mixed *versus* estimated cellular fractions with our NNLS-based deconvolution.

Validation of our deconvolution approach for complex tissue was hampered by the availability of DNAm profiles for samples with a defined composition of cell types. Therefore, we applied our deconvolution to various non-malignant tissue samples from TCGA (Supplemental Figure S4A). Overall, the estimates for the different cell types are compatible with the assumed real cellular composition in the tissue. Furthermore, we have also applied our targeted pyrosequencing approach to various DNA samples of tissues, and the predictions indicated similar composition of different cell types as estimated for the Illumina BeadChip data (Supplemental Figure S4B).

## Discussion

Epigenetic modifications govern cellular differentiation into specific lineages and therefore DNAm is ideally suited for cellular characterization (25-27). Previous approaches for DNAm-based deconvolution of different cell types utilized larger signatures with multiple CpGs of Illumina BeadChip datasets (5, 7, 14). Our proof of concept study demonstrates that estimates for the cellular composition are also feasible by targeted analysis of individual CpGs. There is always a trade-off between different methods: combining a multitude of CpGs into bioinformatic predictors generally increases the precision of epigenetic signatures (28). On the other hand, the precision of DNAm measurements at individual CpGs is higher in pyrosequencing data as compared to β values on Illumina BeadChips (29). Furthermore, microarray analysis is relatively time-consuming and expensive. Currently, classification of cell types is often based on antibody detection of individual epitopes – thus, estimates of the cellular composition by individual CpGs may be feasible, too.

A bottleneck of our analysis is the limited number of defined cellular subsets with available DNAm profiles. The lack of precise measures to distinguish between cell types is also reflected by the ongoing quest of the Human Cell Atlas Project, to define all human cell types in terms of distinctive molecular profiles and to connect this information with classical cellular descriptions (such as location and morphology) (1). For example, fibroblasts and MSCs could possibly resemble the same type of cell (30). On the other hand, fibroblasts are very heterogeneous and can differ greatly depending on their tissue of origin (31, 32). For leukocyte subsets, it has been suggested that particularly cell-subset-specific hypomethylation is permissive for gene expression and regulates corresponding cell functions (33). Indeed, in our analysis cell-type specific CpGs were predominantly hypomethylated – with the exception of iPSCs that resemble a rather non-differentiated ground state. Our hypomethylated CpGs for leukocytes, fibroblasts, endothelial cells, epithelial cells, hepatocytes, glia, neurons and iPSCs cannot span the many facets of cellular classification but at least most cell types can be subordinated to at least one of these broad categories. In previous work, we have extensively studied characteristic DNAm patterns of hematopoietic subsets (16), but we have chosen to not over-represent the hematopoietic compartment in our deconvolution approach and to therefore combine all leukocytes into one category.

Fibroblasts are embedded into the extracellular matrix in native tissue, but there is no distinct cell marker that allows reliable quantification of this subset. Our FibroScore was significantly increased in lung fibrosis and liver cirrhosis. Targeted pyrosequencing of the two CpGs may therefore provide a simple estimate for relative changes of fibroblasts, e.g. for staging of fibrotic diseases. Furthermore, cancer-associated fibroblasts (CAFs) play a central role for tumorigenesis, progression, and metastasis in many cancers (34, 35). It has been shown that the fraction of CAFs, which was estimated for example by the percentage of cells that stained positive for alpha smooth muscle actin, are associated with overall survival in several types of solid cancer (36, 37). Our findings support the notion that an epigenetic fibroblast signature can support stratification of cancer. In the future, it will be important to better understand the epigenetic heterogeneity of CAFs and how these signatures are affected by epigenetic aberrations of the malignant clone.

While the FibroScore may be indicative of the relative fraction of fibroblasts in tissue, it does not provide a quantitative measure for the percentage of fibroblasts. To this end, we have further developed our targeted approach for deconvolution of various cell types in tissue. It is difficult to access the accuracy of our NNLS based deconvolution for tissue samples, since we were lacking precise and validated information on the cellular composition. Nevertheless, the results of the *in-vitro* mixes showed that deconvolution with individual cell-type-specific CpGs is feasible.

## Conclusions

Our results demonstrate that individual CpGs, which are particularly hypomethylated in specific cell-types, can be used to estimate the fraction of fibroblasts or the composition of cellular mixes and tissues. In contrast to genome wide DNAm profiles targeted analysis, e.g. by pyrosequencing, provides new perspectives for small amounts of DNA and to derive robust procedures according to directives for *in vitro* diagnostic devices. Such analysis may be useful to gain insight into the composition of unknown tissue specimen or to correlate the percentage of specific cellular subsets with clinical parameters. Furthermore, it might provide estimates for the composition of cell free DNA (cfDNA), which is increasingly relevant for liquid biopsy (7, 38).

## Methods

### Data acquisition and processing of DNAm profiles

We compiled a curated dataset of DNAm profiles (450K and EPIC Illumina BeadChip platforms) of well characterized and non-malignant human cell types. All analysis and data retrieval was performed with the R programming language v3.6.2 and functions from Bioconductor v3.9. The data was retrieved from Gene Expression Omnibus (through the GEOquery v2.52.0 R package (GEOquery, RRID:SCR_000146), Supplemental Table S1) and data processing was performed using the minfi v1.30.0 R package (minfi, RRID:SCR_012830) and in-house scripts. Features were limited those to CpGs shared between the 450K and EPIC platforms, and we excluded probes related to sexual chromosomes and probes not shared across all samples (missing data), resulting in 415,366 CpGs for further selection. During the data acquisition process (at time of analysis), several samples and features were dropped due to conflicting or missing names, bad file formatting and missing data. In the end we had a total of 579 samples from 14 different cell types. For samples where raw data was available (IDAT files), ssNoob normalization method was applied (39), otherwise no additional normalization of beta values was performed. To avoid bias and overfitting, samples were divided into two independent datasets, a training (n = 409) and validation set (n = 170). Datasets of The Cancer Genome Atlas (TCGA) project (Level 1 methylation array data) were downloaded and preprocessed with the TCGAbiolinks v2.12.6 (TCGAbiolinks, RRID:SCR_017683) and SeSAMe v1.2.0 packages in R (except for TCGA-STAD, which at the time of the analysis was unreachable) following their respective pipelines (Supplemental Table S2) (40, 41).

### Feature selection and signatures for classification and deconvolution

In order to find the best CpGs to perform classification, we subjected the training data to a stratified k-Fold Cross-Validation setup (k=10). We defined *C*_*i*_ as our cell of interest and *C*_*other*_ as the class that englobes all the other cell types. For a given fold, we calculate the difference in means between *C*_*i*_ and *C*_*other*_ (dMean) and the sum of variances within C_i_ and C_other_ (sVar) for each CpG. The relationship between mean and variances is exemplified in Figure 1B, where we assume that CpGs with higher absolute dMean, and lower sVar were considered more discriminative. To capture a set of discriminative CpGs, we define a parabola function and select all CpGs for which *y < (ax)*^*2*^ (Figure 1B). Initially, a is set to 0.1. If less than 10 hypermethylated (x > 0) or 10 hypomethylated (x < 0) CpGs are selected, a is incremented by 0.1 until the previous criteria is reached. Next, we compute the Area Under the Precision-Recall curve (AUPR) on the remaining folds and scale it by the absolute dMean (42). We consider here hypo- and hypermethylated CpGs separately for estimating dMean. The above procedure is repeated for each fold. A final score is obtained by the average scaled AUPR multiplied by the proportion of folds where a CpG was selected as a top scoring candidate. This is then used to obtain a final CpG ranking. This measure selects CpGs present in more folds, having higher AUPR and higher absolute dMean. The best iPSC CpG was not suitable for primer design for pyrosequencing and therefore the second best was selected for this cell type. For the FibroScore, we used the F1-score (without scaling) and selected from the best CpGs, one hypo- and one hypermethylated CpG, after initial screening with pyrosequencing.

### Deconvolution of cell type proportions

Using the cell type specific CpGs previously selected for classification and their mean methylation value (for each cell type) on the training dataset as our reference matrix, we applied a reference-based non-negative least-squares (NNLS) algorithm (16, 23). An application for cell type deconvolution is provided as separate Excel tool (Supplemental Table S3) and as the DeconvolutionApp, https://costalab.ukaachen.de/shiny/tmaie/deconapp/ (accessed 24^th^ July 2020).

### Survival analysis

Survival analysis and plots, using the median FibroScore as the stratification factor, were performed on TCGA data (Supplemental Table S2) with the survival v3.1.12 and survminer v0.4.6 packages in R (43, 44). P-values are based on the log-rank test.

### Cell Culture

Human mesenchymal stromal cells (45), dermal fibroblasts (32), human umbilical vein endothelial cells (HUVECs) (46), and iPSCs (45, 47) were isolated and thoroughly characterized as described in our previous work. Human cell lines HepG2, HuH-7, Hep3B, and HaCat were maintained at RWTH Aachen Medical School under standard culture for isolation of genomic DNA. For HepG2 and HaCat DNA was directly isolated from cryopreserved vials.

### Isolation of genomic DNA and bisulfite conversion

Genomic DNA from cells and tissues was isolated with the NucleoSpin® Tissue Kit (Macherey-Nagel), and from blood (150 µl) with the QIAamp DNA Blood Mini Kit (Qiagen). DNA concentration was measured using the NanoDrop™ 2000 spectrophotometer (Thermo Scientific™) and bisulfite converted using the EZ DNA Methylation Kit (Zymo Research).

### Pyrosequencing

Bisulfite converted DNA was amplified with a region-specific biotinylated/unmodified DNA primer pair (Metabion; Supplemental Table S4) using the PyroMark PCR Kit (Qiagen) according to the manufactures instructions: Initial activation at 95°C for 15 min, then 45 cycles of 30 s at 94°C, 30 s at 56°C and 30 s at 72°C followed by a final extension at 72°C for 10 min. Pyrosequencing was performed on the PyroMark Q96 and the Q48 Autoprep platforms. The results were analyzed using the Pyro Q-CpG 1.0.9 or the PyroMark Q48 Advanced Software, respectively.

### Quantification and statistical analysis

In total we used DNAm profiles of 579 samples from 46 different studies for training and validation sets. The pyrosequencing signatures were validated with four cell lines, 12 MSC samples, 6 fibroblast samples, 4 HUVEC preparations, 5 iPSC lines, 8 blood samples and 14 different tissue samples. To estimate the significance of differential DNAm and FibroScore in the lung fibrosis and liver cirrhosis datasets we utilized the two-sided t-test: *** < 0.001, ** < 0.01, * < 0.05. P-values for overall survival in cancer are based on the log-rank test.

## Declarations

### Ethics approval and consent to participate

Whole blood and tissue samples were taken after informed consent according to the guideline of the local ethics committee (EK 206/09) and provided by the RWTH Centralized Biomaterial Bank (RWTH cBMB).

### Authors’ contributions

Conception of the project: W.W. and I.C.; curation of datasets: M.S. and T.M.; analysis of DNAm profiles: T.M. and M.S.; Pyrosequencing: M.S.; Support for cell and tissue preparations: E.D.; the initial draft of this manuscript was written by M.S. and reviewed and edited by all authors.

## Acknowledgements

Tissue samples used in this study were provided by the RWTH centralized biobank (RWTH cBMB) of the Medical Faculty of RWTH Aachen Medical School.

