## Supplemental Figures S1-S4 and Tables S1, S2 and S4 for "Deconvolution of cellular subsets in human tissue based on targeted DNA methylation analysis at individual CpG sites"

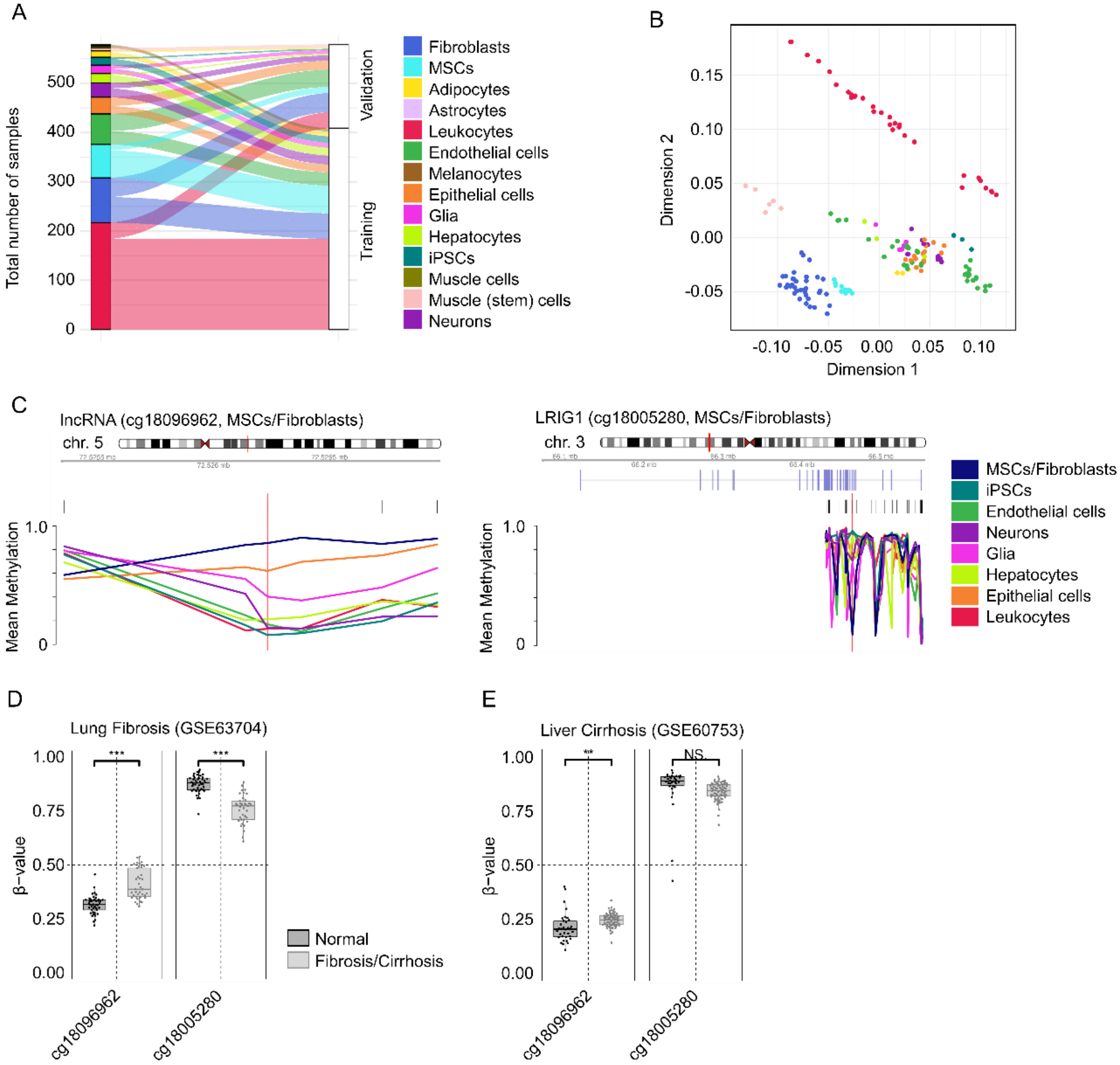


**Supplemental Figure S1: Selection of cell-type specific CpGs for fibroblasts**

(A) Split of total samples (n = 579) into training (n = 409) and validation set (n = 170).

(B) Multidimensional scaling (MDS) plot of the validation data (n = 170) shows that samples cluster by cell type. For the analysis, all CpGs shared between the 450K and the EPIC BeadChip were included (except XY chromosomes).

(C) Chromosomal location and association with corresponding genes for the selected fibroblast-specific CpGs: lncRNA (*RP11-60A8.1*) and leucine rich repeats and immunoglobulin like domains 1 (*LRIG1*). Shown are mean β values from the training data set of neighboring CpGs for the respective cell types. The red lines depict the relevant CpGs.

(D) DNAm levels (β values) of the two selected CpGs from the FibroScore in the lung fibrosis dataset GSE63704 (450K BeadChip) (1). Two-sided t-test: *** $p < 0.001$.

(E) DNAm levels (β values) of the two selected CpGs from the FibroScore in the liver cirrhosis dataset GSE60753 (450K BeadChip) (2). Two-sided t-test: : ** $p < 0.01$, NS = not significant.

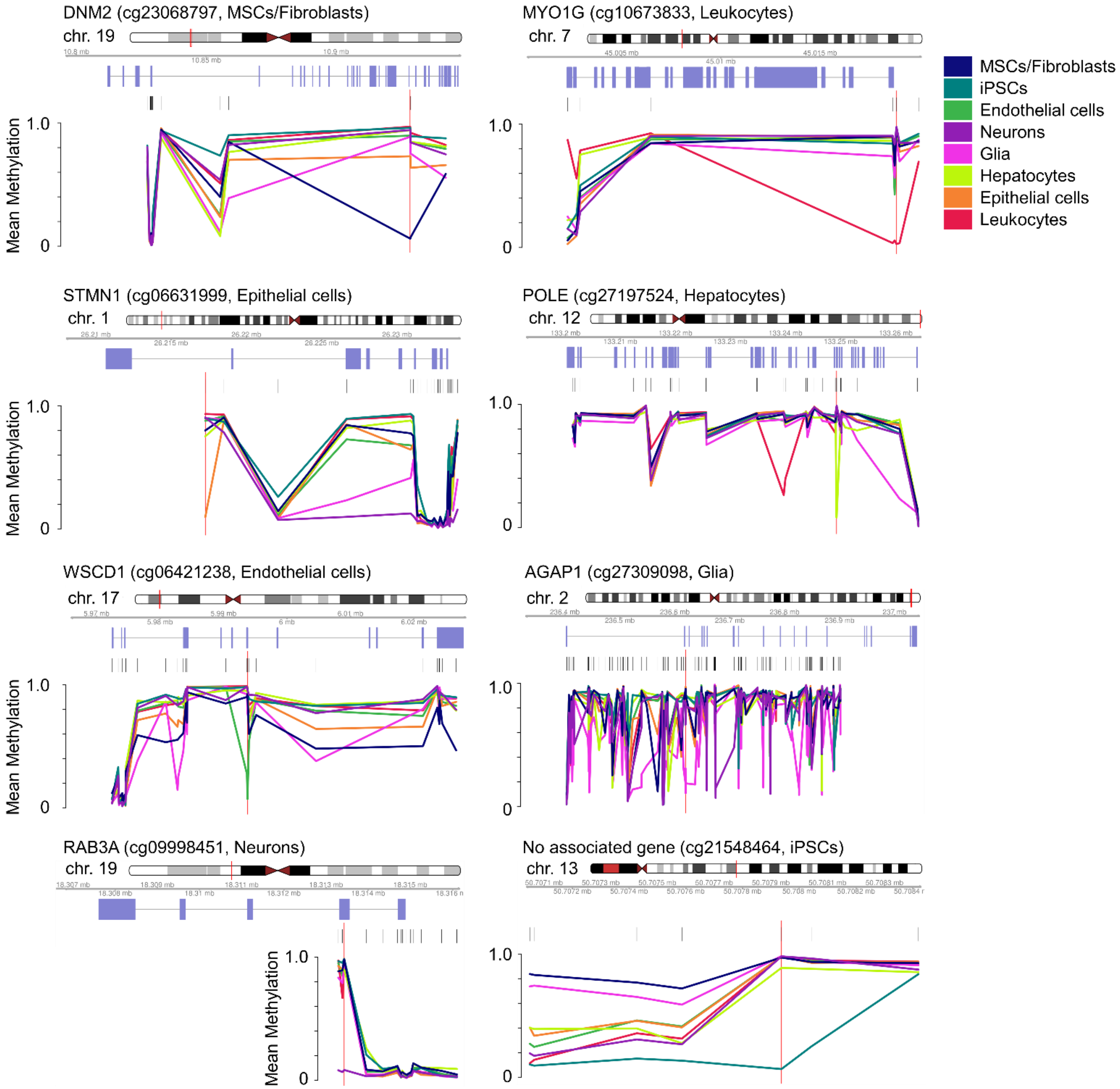


**Supplemental Figure S2: Genomic context of cell-type-specific CpG sites**

Chromosomal location and association with corresponding genes for the selected cell-type-specific CpGs: Dynamin 2 (*DNM2*), Myosin IG (*MYO1G*) encodes for the human minor histocompatibility antigen HA-2, which is only expressed in hematopoietic cells (3), Stathmin 1 (*STMN1*), DNA polymerase epsilon catalytic subunit A (*POLE*), WSC Domain Containing 1 (*WSCD1*), ArfGAP With GTPase Domain, Ankyrin Repeat And PH Domain 1 (*AGAP1*) is associated with neurodevelopmental disorders and overexpressed in brain (4), RAS-Associated Protein RAB3A (*RAB3A*) is also highly expressed in brain (4). Shown are mean β values from the training data set of neighboring CpGs for the respective cell types. The red lines depict the relevant CpGs.

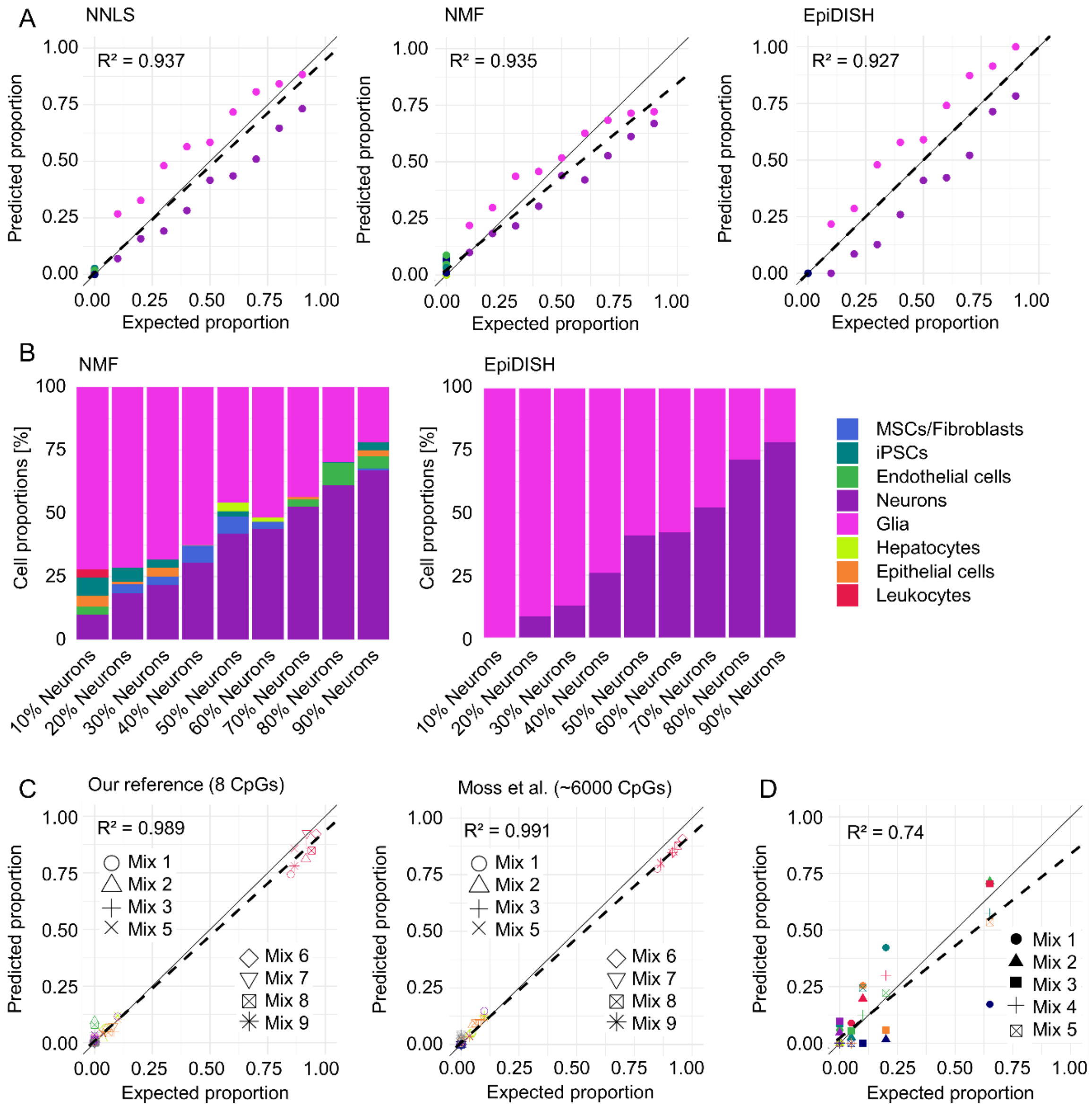


**Supplemental Figure S3: Comparison of different deconvolution methods**

(A) In addition to the NNLS model, two alternative deconvolution approaches were considered for *in vitro* neuron-glia-DNA mixes from dataset GSE41826 (5): non-negative matrix factorization (NMF) and EpiDISH (6, 7). Comparison of expected *versus* predicted DNA proportions by deconvolution with different algorithms (NNLS, NMF and EpiDISH) (6, 7).

(B) The predicted cell fractions for the neuron-glia-DNA mixes from dataset GSE41826 (5) are depicted based on the eight cell-type-specific CpGs for NMF and EpiDISH. The best correlation of the predicted cell fractions with the real mixture of neurons/glia was observed for the NNLS-based deconvolution (Figure 4), while EpiDISH did not misclassify any other cell type.

(C) Cell type DNA mixes from dataset GSE122126 (8). Comparison of expected *versus* predicted DNA proportions by deconvolution with different reference matrices.

(D) Pyrosequencing of DNA mixes. Comparison of expected *versus* predicted DNA proportions by deconvolution.

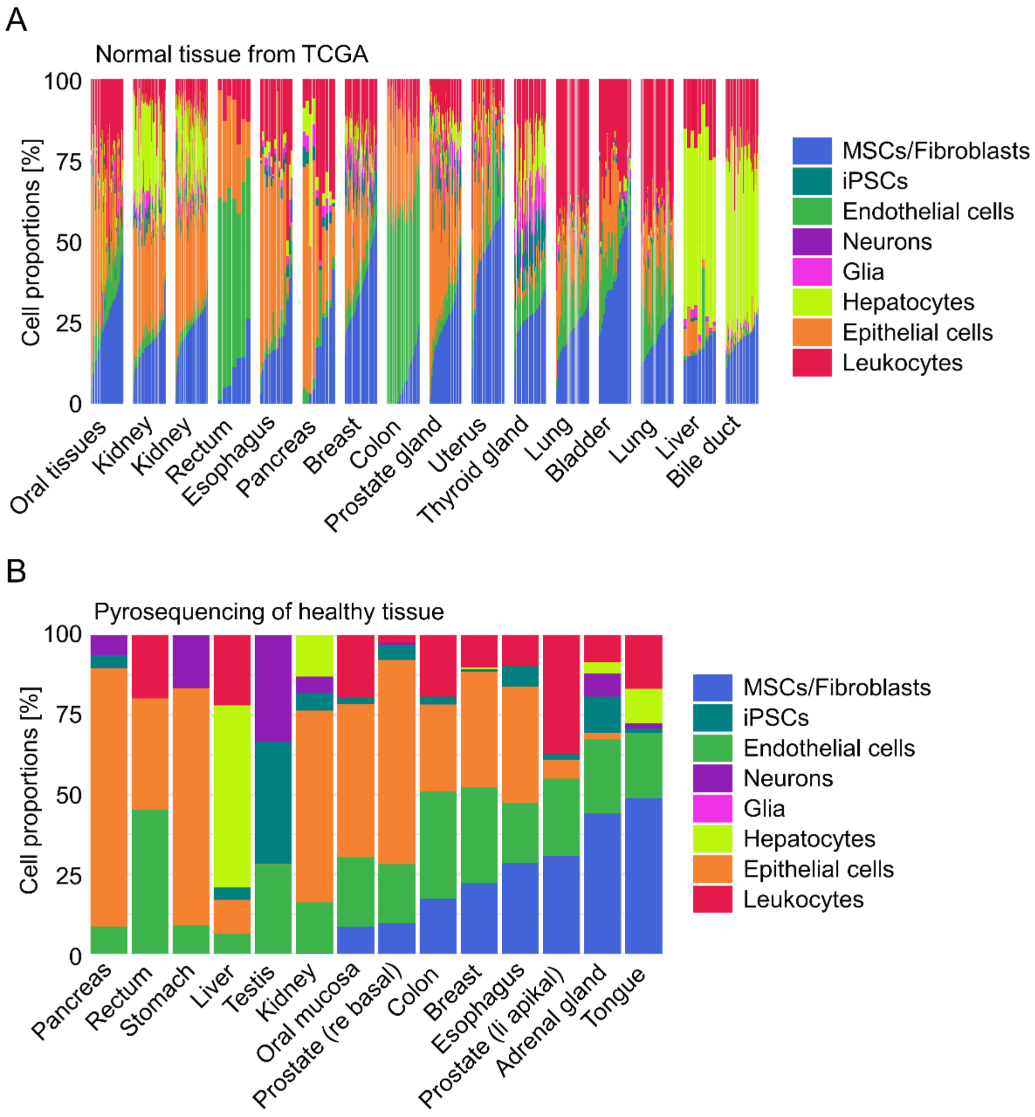


**Supplemental Figure S4: Deconvolution of cell mixtures based on individual cell-type-specific CpGs**

(A) Deconvolution of normal tissues from the TCGA database (450K BeadChip). Shown are estimated cellular fractions. The relevant CpG for neurons was not available for TCGA datasets and therefore left out.

(B) Deconvolution of healthy tissues based on pyrosequencing of DNAm at the eight relevant CpGs.

**Supplemental Table S1. 450k/EPIC Illumina BeadChip datasets used in this study**

| GSE | SAMPLE TYPE | NUMBER OF SAMPLES | REFERENCES |
|---|---|---|---|
| **Training dataset** | | | |
| GSE34486* | endothelial cells | 10 | (9) |
| GSE40699 | muscle cells, epithelial cells, hepatocytes, fibroblasts, astrocyte, endothelial cells | 26 | (10) |
| GSE41933 | mesenchymal stromal cells | 12 | (11) |
| GSE43976 | leukocytes | 10 | (12) |
| GSE50222 | leukocytes | 8 | (13) |
| GSE52025 | fibroblasts | 2 | (14) |
| GSE52112 | mesenchymal stromal cells | 34 | (15) |
| GSE58622 | adipocytes | 10 | (16) |
| GSE59065 | leukocytes | 20 | (17) |
| GSE59091 | induced pluripotent stem cells, fibroblasts | 12 | (18) |
| GSE59250 | leukocytes | 60 | (19) |
| GSE59796 | leukocytes | 4 | (20) |
| GSE60753 | hepatocytes | 15 | (2) |
| GSE63409 | leukocytes | 30 | (21) |
| GSE65078 | induced pluripotent stem cells, fibroblasts | 8 | (22) |
| GSE68134 | induced pluripotent stem cells, fibroblasts | 6 | (23) |
| GSE71955 | leukocytes | 20 | (24) |
| GSE74877 | epithelial cells, melanocytes, mesenchymal stromal cells, fibroblasts, endothelial cells | 9 | (25) |
| GSE77135 | fibroblasts | 20 | (26) |
| GSE79144 | glia, neurons | 18 | (27) |
| GSE79695 | mesenchymal stromal cells | 12 | (28) |
| GSE82234 | endothelial cells | 3 | (29) |
| GSE85647 | leukocytes | 6 | (30) |
| GSE87095 | leukocytes | 10 | (31) |
| GSE87177 | endothelial cells | 1 | (32) |
| GSE88824 | leukocytes | 16 | (33) |
| GSE92843 | epithelial cells | 1 | (34) |
| GSE95096 | fibroblasts | 1 | (35) |
| GSE98203 | neurons | 10 | (36) |
| GSE99716 | endothelial cells | 1 | (37) |
| GSE103253* | endothelial cells | 10 | (38) |
| GSE107226 | fibroblasts | 4 | (39) |
| **Validation dataset** | | | |
| GSE34486* | endothelial cells | 6 | (9) |
| GSE51921 | iPSCs, fibroblasts | 6 | (40) |
| GSE53302 | muscle stem cells | 6 | (41) |
| GSE68851 | fibroblasts | 12 | (42) |
| GSE71244 | leukocytes | 24 | (43) |
| GSE74486 | glia, neurons | 15 | (44) |
| GSE85566 | epithelial cells | 6 | (45) |
| GSE86258 | fibroblasts | 7 | (46) |
| GSE86829 | fibroblasts | 3 | (47) |
| GSE87797 | mesenchymal stromal cells | 12 | (48) |
| GSE103253* | endothelial cells | 15 | (38) |
| GSE104287 | leukocytes | 8 | (49) |
| GSE106099 | feto-placental endothelial cells | 12 | (50) |
| GSE109042 | buccal epithelial cells | 6 | (51) |
| GSE111396 | fibroblasts | 14 | (52) |
| GSE122126 | adipocytes, neurons, hepatocytes, endothelial cells, epithelial cells, leukocytes | 18 (13 EPIC) | (8) |
| **Other datasets** | | | |
| GSE41826 | glia-neuron DNA mixes | 9 | (5) |
| GSE60753 | normal and cirrhotic liver | 100 | (2) |
| GSE63704 | normal and fibrotic lung | 80 | (1) |
| GSE122126 | cell type DNA mixes | 8 | (8) |

* Samples from these studies were considered either for training or validation sets

**Supplemental Table S2. Association of fibroblast-associated DNAm with overall survival in cancer**

| TCGA PROJECT | CANCER TYPE | SURV_PVAL CG18096962 | SURV_PVAL CG18005280 | SURV_PVAL FIBROSCORE |
|---|---|---|---|---|
| TCGA-ACC | Adrenocortical Carcinoma | 0.0001* | 0.7525 | 0.0003* |
| TCGA-MESO | Mesothelioma | 0.2466 | 0.0001* | 0.0011* |
| TCGA-HNSC | Head and Neck Squamous Cell Carcinoma | 0.7474 | 0.0153* | 0.0050* |
| TCGA-PCPG | Pheochromocytoma and Paraganglioma | 0.1130 | 0.9490 | 0.0064* |
| TCGA-KICH | Kidney Chromophobe | 0.0170* | 0.1673 | 0.0119* |
| TCGA-KIRC | Kidney Renal Clear Cell Carcinoma | 0.2481 | 0.0103* | 0.0619 |
| TCGA-BLCA | Bladder Urothelial Carcinoma | 0.3357 | 0.2406 | 0.0623 |
| TCGA-THYM | Thymoma | 0.1658 | 0.1807 | 0.1081 |
| TCGA-PRAD | Prostate Adenocarcinoma | 0.7707 | 0.9890 | 0.1087 |
| TCGA-SARC | Sarcoma | 0.1208 | 0.3432 | 0.1390 |
| TCGA-UCEC | Uterine Corpus Endometrial Carcinoma | 0.6565 | 0.7381 | 0.1453 |
| TCGA-LGG | Brain Lower Grade Glioma | 0.0113* | 0.0177* | 0.1704 |
| TCGA-THCA | Thyroid Carcinoma | 0.1991 | 0.5390 | 0.2141 |
| TCGA-LUAD | Lung Adenocarcinoma | 0.4578 | 0.3120 | 0.2541 |
| TCGA-BRCA | Breast Invasive Carcinoma | 0.4685 | 0.8774 | 0.2661 |
| TCGA-ESCA | Esophageal Carcinoma | 0.7665 | 0.4520 | 0.3506 |
| TCGA-READ | Rectum Adenocarcinoma | 0.6664 | 0.2644 | 0.4358 |
| TCGA-LIHC | Liver Hepatocellular Carcinoma | 0.5432 | 0.8678 | 0.4505 |
| TCGA-PAAD | Pancreatic Adenocarcinoma | 0.9590 | 0.0090* | 0.4868 |
| TCGA-TGCT | Testicular Germ Cell Tumors | 0.5297 | 0.7229 | 0.4901 |
| TCGA-LAML | Acute Myeloid Leukemia | 0.2399 | 0.9467 | 0.4939 |
| TCGA-CHOL | Cholangiocarcinoma | 0.2521 | 0.2828 | 0.5361 |
| TCGA-DLBC | Lymphoid Neoplasm Diffuse Large B-cell Lymphoma | 0.8924 | 0.4410 | 0.5602 |
| TCGA-UCS | Uterine Carcinosarcoma | 0.0179* | 0.0865 | 0.6019 |
| TCGA-GBM | Glioblastoma Multiforme | 0.7417 | 0.3277 | 0.6096 |
| TCGA-SKCM | Skin Cutaneous Melanoma | 0.6751 | 0.7819 | 0.6126 |
| TCGA-UVM | Uveal Melanoma | 0.3799 | 0.0001* | 0.6207 |
| TCGA-CESC | Cervical Squamous Cell Carcinoma & Endocervical Adenocarcinoma | 0.5976 | 0.3391 | 0.6233 |
| TCGA-OV | Ovarian Serous Cystadenocarcinoma | 0.6395 | 0.0922 | 0.6395 |
| TCGA-COAD | Colon Adenocarcinoma | 0.7778 | 0.7443 | 0.7918 |
| TCGA-LUSC | Lung Squamous Cell Carcinoma | 0.9646 | 0.4866 | 0.9541 |
| TCGA-KIRP | Kidney Renal Papillary Cell Carcinoma | 0.2736 | 0.0455* | 0.9929 |

* $p < 0.05$

**Supplemental Table S3. Application for cell type deconvolution**

An application for cell type deconvolution is provided as separate Excel tool. The mean DNAm values for the cell-type-specific CpGs of the Illumina BeadChip training dataset are given as reference matrix. Furthermore, the application allows NNLS predictions to estimate the cellular composition in independent datasets. This table was generated in analogy to the NNLS application for Epi-Blood-Count (53).

**Supplemental Table S4. Primer DNA sequences used for pyrosequencing**

| NAME | DNA SEQUENCE |
|---|---|
| cg18096962_Forward | GAGTATTGGGTTTATTTAGTTTTAGGAT |
| cg18096962_Reverse_Biotin | TCAAATTCTATTTACTACCCTCTTCC |
| cg18096962_Sequencing | GTTTTTATTTTTGAG |
| cg18005280_Forward | TATTGGTGTTATTGGGGGAGG |
| cg18005280_Reverse_Biotin | CCCACAACCATTCTAAAACAATC |
| cg18005280_Sequencing | TGTTATTGGGGGAGG |
| cg10673833_Forward | TGTTGTTAGGGTTGGAAGTTAATTT |
| cg10673833_Reverse_Biotin | CACCAACCTCCTCCAATACTAATATAA |
| cg10673833_Sequencing | GGGGAGGATTTAGT |
| cg06421238_Forward | TTGTGGGGGATGGGTAGT |
| cg06421238_Reverse_Biotin | ACCTCCTCCCTACAAATCCTATATCT |
| cg06421238_Sequencing | GATAAAGTTTAGGAAGAGGTT |
| cg06631999_Forward_Biotin | GGGTTGTTTTTTGGTTATTAGAGTTAGGTA |
| cg06631999_Reverse | CTCTTCTTTCCAATTTTTTCCAAATAATC |
| cg06631999_Sequencing | CTCTAACTCAATCCCTAAATAC |
| cg23068797_Forward_Biotin | TTTTTGGGTTTAGGAGGAATGTT |
| cg23068797_Reverse | CCAACTAATACCACATCTAAAACTATTTACAATAC |
| cg23068797_Sequencing | AATACCACATCTAAAACTATTTAC |
| cg27309098_Forward_Biotin | GAGGAAATTGAGGTTTAGAGATATGAA |
| cg27309098_Reverse | CTAAAATCAAACTTAAAATACAACTCCTTAATA |
| cg27309098_Sequencing | CAAAAAAATTTTTACC |
| cg27197524_Forward_Biotin | GAAGAATTTGAATTTTAGGGAAGAAGTAT |
| cg27197524_Reverse | CCCAACAAACAAAAACAAAAATTCAATA |
| cg27197524_Sequencing | TACTAAACTCTAAAACCTAC |
| cg21548464_Forward | GTTTTTTAGTTGGGATTTATTTAGATTTGT |
| cg21548464_Reverse_Biotin | AACCCTTACCATCTTCTACCTAAACT |
| cg21548464_Sequencing | CATATTCAAAATTCTCATCAT |
| cg09998451_Forward_Biotin | GGGAATTTTGTATTTTAGTTGTGGATTTTT |
| cg09998451_Reverse | TAAACCTCAATTAACCCCTACTCAA |
| cg09998451_Sequencing | GTAAAATTTTGTTTGAT |
